# A positive statistical benchmark to assess network agreement

**DOI:** 10.1101/2022.10.21.513307

**Authors:** Bingjie Hao, István A. Kovács

## Abstract

As the current best practice, an experimental network dataset is validated by showing significant overlap with a gold standard network. Significance is assessed by comparison to a negative benchmark, often a randomized version of the same gold standard. While such analysis can reliably indicate the presence of signal, it is ill-suited to assess how much signal there is. As an alternative, here we introduce a positive statistical benchmark corresponding to the best-case scenario, capturing the maximum possible overlap between two networks. Such a positive benchmark can be efficiently generated in a maximum entropy framework and opens the way to assess if the observed overlap is significantly different from the best-case scenario. In combination with the negative benchmark, we provide a normalized overlap score (*Normlap*). As an application, we compare molecular and functional networks, resulting in an *agreement network* of human as well as yeast network datasets. Although the number of shared interactions between most networks within the same organism is relatively small, we find that it is typically close to the best-case scenario. This paradox is resolved by the underlying *degree inconsistency*, meaning that highly connected hubs in one network often have small degrees in another, limiting the potential overlap. Furthermore, we illustrate how Normlap improves the quality assessment of experimental networks, fostering the creation of future high-quality networks.

## 1 Introduction

We are witnessing a rapid expansion of network data, transforming a broad range of scientific fields^1^. In network representations, entities of interest are represented by nodes and connected by links. Real-life networks often have a broad degree distribution, where node degree denotes the number of links connected to each node^2^. In social and biological applications, multiple network maps exist that capture different relationships between the same nodes. Even against recent advances, it remains fundamentally impossible to obtain complete maps of macromolecular networks using only a single assay^3^. For example, various biophysical relationships between proteins are critical to biological processes in cells.

There are two major categories of the corresponding experimental assays, *binary methods* such as Yeast-Two-Hybrid (Y2H) and protein-fragment complementation assays (PCA), which require direct contact of the proteins (protein-protein interactions, PPIs)^4–8^ or *non-binary methods*, such as affinity purification followed by mass-spectrometry (AP-MS)^9–12^, where interacting proteins are not necessarily in direct contact. Such biophysical measurements have provided a wealth of overlapping, yet distinct experimental data about both binary and non-binary PPIs. In addition, functional networks such as genetic interaction (GI) networks^13,14^ identify functional relationships among the genes or their corresponding gene products. Quality control is essential to utilize any of these networks for a comprehensive understanding of the underlying biological processes.

Unfortunately, reliable quality control of network maps remains a challenging open question. Traditionally, the quality of network datasets is evaluated by counting the overlap either computationally with another, gold standard network or experimentally with a complementary assay^3–5,15^. However, such a comparison often leads to a disturbingly low overlap^3^. For example, in the systematic genome-wide human binary protein interactome (HuRI)^5^, among 52,569 PPIs observed in three experimental Y2H assays, only 2,707 (5%) PPIs are supported by at least two assays, while only 258 (0.5%) are found in all three assays. The question naturally arises: Is such a low overlap an indication of low network quality?

Previous studies have shown that even if all measured interactions are true positives, the low observed overlap could stem from multiple sources^15^. First of all, current biological networks are still highly incomplete^5,6,16^. Taking the most studied organism – yeast – as an example, a combination of PPIs from four systematic Y2H assays covered only 12 − 25% of the estimated yeast binary interactome^6^. Hence, a low overlap could easily arise from a high level of incompleteness, especially if the data coverage is more homogeneous in some datasets than in others^17^. As an additional factor, experimental assays are often designed to be complementary to each other to unveil as much new information as possible^3^. Overall, different assays will preferably capture some interactions versus others^3,6,17^, leading to the detection of distinct sets of links and thus low overlap between the various datasets. As a result of these factors, the observed node degrees can also be inconsistent across network maps, even if the degree distributions appear to be similar. For example, in Fig. 1a-b, a highly connected node, or ‘hub’, in one network may have a small node degree in the other network and vice versa, causing the degree correlation plot to be off diagonal and hindering a significant overlap. The problem goes beyond the extent to which the reported node degrees can be trusted. Such node degree inconsistency questions the validity of using the observed overlap as a measure of network quality, since a low overlap could either come from degree inconsistency or quality issues. The use of overlap to assess network quality is especially problematic when comparing different types of networks, where the degree distribution could be wildly different and some networks might not even have highly connected hubs. The example in Fig. 3a shows that binary physical networks showcase a broad degree distribution, while functional networks such as GI networks tend to have a narrow distribution, with no major hubs. Undoubtedly, such a qualitatively different degree distribution hinders a consistent degree sequence. Thus, node degrees need to be considered as confounding factors, just like in state-of-the-art methods to generate negative benchmarks (null model). In the null models, network randomization is performed while preserving node degrees either exactly or on average, as a soft constraint, to compare network characteristics^18,19^. Although a significant difference from the negative benchmark indicates the presence of signal in the network of interest, it fails to quantify the amount of signal.

**Figure 1.**
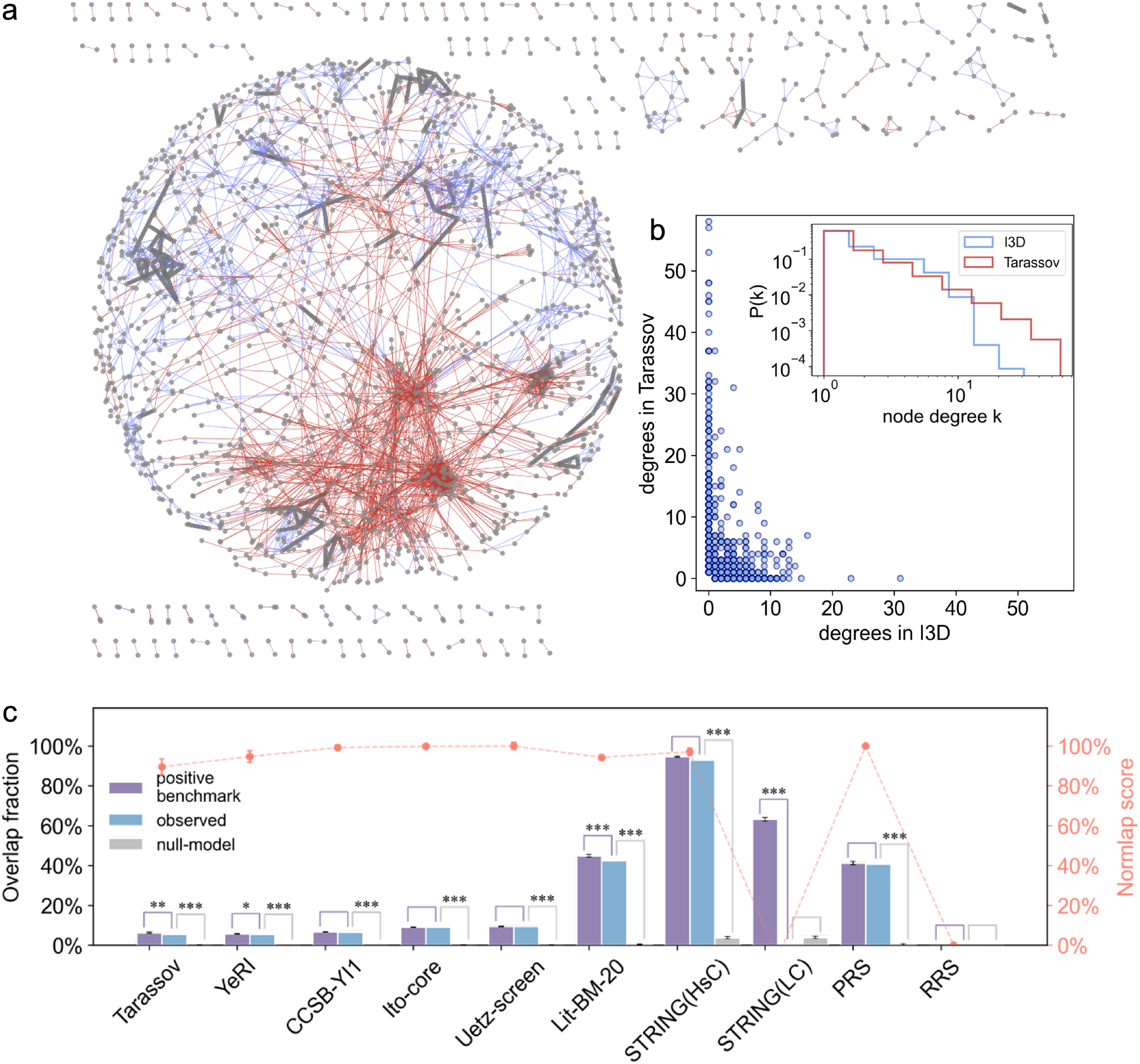
Yeast PPI networks compared to the gold standard network I3D and the null model. (a) The network structure of I3D (blue) and Tarassov (red). The 137 overlapping links (2%) are colored in dark gray. (b) Degree mismatch between I3D and Tarassov networks, with the degree distributions shown in the inset. (c) The overlap fraction of various PPI networks compared to I3D. The proposed positive benchmark is shown in purple, leading to the normalized overlap (Normlap) score in red. In gray, we show a degree-preserved randomized I3D as a null model for reference. The significance of the observed overlap compared to the null model or the positive benchmark is indicated by *: *p* < 0.05, **: *p* < 0.005, ***: *p* < 0.0005.

Here, as the so far missing another side of the assessment, we set out to establish a positive statistical benchmark to compare against, as a best-case scenario. Generally, in addition to preserving node degrees, a generative model is needed to compare any observed variable with a benchmark, through actual instances of the benchmark ensemble. In the proposed positive benchmark, every link in both networks is assumed to be sampled from the same underlying network of real connections, according to the observed degrees. Such a positive benchmark corresponds to a best-case scenario, where the two networks are the same as far as the node degrees allow. The overlap between the two sampled networks in the best-case scenario is regarded as the *positive benchmark* for the network overlap. With both the negative and positive benchmarks at hand, we can then place the observed overlap on a scale between 0 (null-model) and 100% (best-case scenario), leading to the normalized network overlap or ‘Normlap’ score.

Due to the availability of a broad range of molecular and functional network datasets in yeast, we selected *S. cerevisiae* as the primary model system, in addition to a smaller cohort of human networks, as listed in the Methods section. In both organisms, we create an agreement network of networks, illustrating the knowledge landscape of the available datasets. In addition, we illustrate how the Normlap score can provide a computational alternative to validate experimental network maps^20,21^.

## 2 Results

### Using a negative benchmark

The negative benchmark is obtained by taking the overlap between the network of interest and the degree-preserved randomized gold standard network. Here, we construct a negative benchmark using a maximum-entropy framework^19,22^ (see Methods for details). As a starting point, we compare the observed overlap of ten yeast PPI networks with a gold standard network I3D and its randomized version as a null model. All networks show significantly larger observed overlap with I3D compared to the null-model (*p* < 0.0005) except for the random reference dataset (RRS)^4^ and a low confidence (LC) PPI network compiled from the STRING database^23^, as shown in Fig. 1c. For simplicity, we will say that we have signal if the observed overlap is significantly higher (*p* < 0.05) than the negative benchmark. Besides, all systematic networks (Tarassov, YeRI, CCSB-YI1, Ito-core and Uetz-screen) have significantly lower overlap (< 10%) than the literature-curated network Lit-BM-20 and the highest confidence (HsC) network from STRING, as well as the positive reference dataset PRS^4^. This observation is in line with the more homogeneous coverage of the interactome by systematic networks compared to integrated networks, as shown previously in Ref. [6]. As an underlying reason for the low overlap fraction, degree inconsistency is commonly observed between these networks, as illustrated for two networks of similar size, I3D and Tarassov in Fig. 1a,b. Hubs in one network have small degrees in the other network and vice versa. The degree inconsistency becomes even more pronounced when we compare networks of different sizes or types such as PPI networks versus GI networks as shown in Fig. 3a. Although a significant overlap relative to the null model indicates the presence of signal in the network of interest, the small observed overlap fraction resulting from the degree inconsistency makes it appear that most of these networks could be problematic. To resolve this issue, we need to establish a quantitative scale where the observed overlap appears between the worst-case and the best-case scenarios.

### Generating a positive statistical benchmark

A straightforward way to normalize the observed overlap between two networks is to divide the overlap by the number of links in the smaller network^24^. This method is overly optimistic as it takes the size of the smaller network as an upper bound, corresponding to the assumption that all links in the smaller network can be contained by the other network. As an alternative, we first introduce a ‘naïve’ upper bound described as follows. The maximum possible overlap is determined by summing up the minimum degrees of each node in the two networks. We then divide the sum by two, as each potential link within a network is considered twice from the perspective of the two nodes it connects. This naïve upper bound typically yields a much lower reference overlap than the size of the smaller network, while still being an exact upper bound, as illustrated in Fig. S1a. However, even the naïve upper bound is not always achievable since it only considers degree constraints independently for each node instead of all degree constraints simultaneously. Thus, a *generative* model is needed to statistically compare the observed overlap with the upper bound.

Hence, next, we propose a generative positive benchmark that takes both the degree inconsistency and the network structure into account. The intuition behind our positive benchmark can be illustrated in the example of mixing salt and pepper. Imagine that we mix two bottles (initial networks) of salt and then split them into two new bottles. There will be a perfect agreement between the content of the two new bottles and they will be identical to the original bottles. However, if one of the bottles is replaced by pepper instead, when we mix up and split them, the new bottles will contain a mixture that is different from the initial bottles. In this case, the new bottles will have a much higher agreement with each other than the original bottles, indicating that the original bottles were indeed very different. Analogously, we take the union of two networks and split the union randomly into two alternative networks according to the observed degree sequences. The overlap between the two alternative networks forms the positive benchmark, corresponding to the best-case scenario. This way, we generate actual random instances of the best-case scenario with specific overlapping links, with the overlap between the two alternative networks serving as the positive benchmark. The proposed positive benchmark preserves the degree sequence (on average) during the randomization process (see Methods for more details), keeping a major factor in control that impacts the network overlap. If the two input networks are really sampled from the union, this process builds a statistical ensemble close to the original level of overlap. If, however, we sample from a complete graph instead of the union of the two networks, i.e., perform degree-preserved network randomization, we arrive at our negative benchmark. In practice, it is sufficient to randomize only the gold standard network for the positive and negative benchmarks (one-sided statistics, Fig. 2a). In network comparisons without a clear gold standard, we randomize one of the networks and calculate the negative and positive benchmarks respectively. The negative (positive) benchmark that is closer to the observed overlap is used as the final negative (positive) benchmark. As expected, the overlap with instances of the positive benchmark are always equal to or lower than the naïve upper bounds with the same degree sequences (Fig. S1a), while also being realizable.

**Figure 2.**
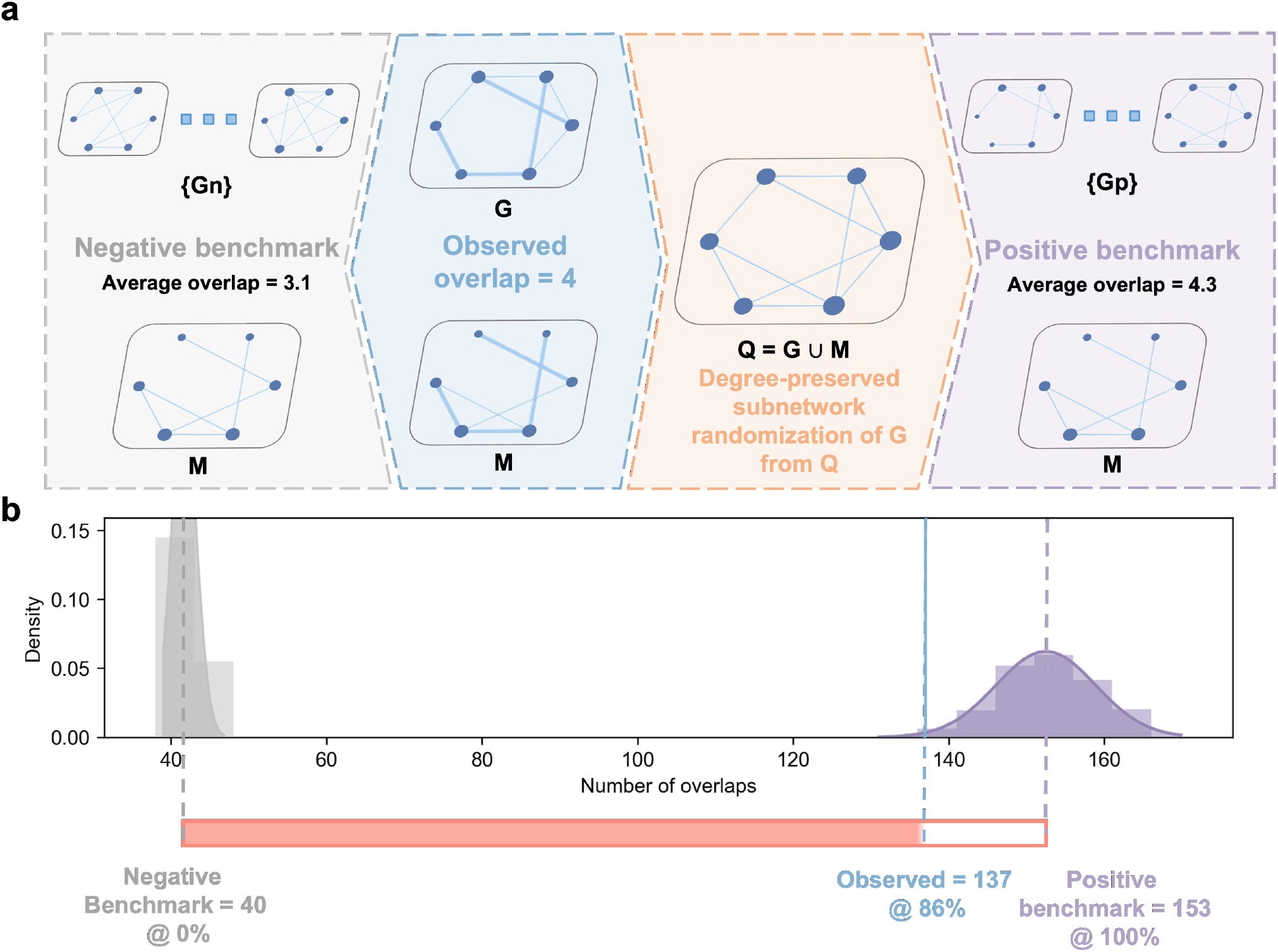
Benchmarks and normalization of network overlap. (a) The negative benchmark is obtained by counting the overlap between the network of interest *M* and the degree-preserved randomized gold standard networks {*Gn*}. The positive benchmark is the overlap between *M* and {*Gp*}, where {*Gp*} are obtained by randomly selecting links from the union *Q* of the gold standard network *G* and *M* according to the degree sequence of *G*. Thicker links indicate the observed overlap. (b) Example of normalizing the observed overlap (blue) with the negative benchmark (gray) and the positive benchmark (purple) between the gold standard network I3D (*G*) and the network of interest, Tarassov (*M*).

Indeed, just like testing if the observed overlap is significantly different from the null model, having access to the positive benchmark allows us to check if the observed overlap is significantly different from the best-case scenario. As illustrated in Fig. 1c, the observed overlaps of CCSB-YI1, Ito-core, Uetz-screen, STRING(HsC) and PRS are not significantly different from the positive benchmark, meaning that these networks are consistent with I3D apart from the observed degree inconsistency. For simplicity, we will call two networks *compatible* if there is signal and the observed overlap is not significantly lower than the positive benchmark. In contrast, if the observed overlap is significantly lower than the positive benchmark, it means the low observed overlap cannot be explained by the degree inconsistency alone and may indicate the presence of substantially different biological mechanisms or even false positives. For instance, the observed overlap between I3D and STRING(LC) is significantly lower (*p* < 0.005) than the positive benchmark because STRING(LC) consists of low-confidence interactions, some of which are likely false positives.

### Normalized network overlap

The positive benchmark opens the way to normalize the observed overlap in combination with the negative benchmark, leading to

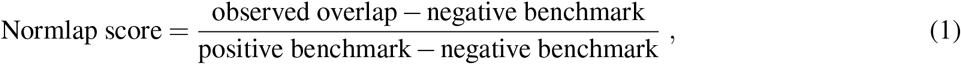

where the *observed overlap* is the number of overlapping links between the network of interest *M* and the gold standard network *G*. The Normlap score places the observed overlap in the range spanned by the negative and positive benchmarks, respectively. Fig. 2b shows an example of normalizing the observed overlap for the Tarassov network with I3D as gold standard.

A major limitation of the observed overlap is its sensitivity to data incompleteness. Thus, we explored if the Normlap score is more robust against incompleteness, potentially providing a less biased view of data quality than the observed overlap. As the most suitable experimental system to test this idea, we considered a systematic proteome-wide human PPI network, HuRI^5^, constructed from nine screens, three for each of three different experimental assays. We compared the network of interest, i.e. the network complied from a number of selected screens from HuRI, with Lit-BM-17^5^ as the gold standard. As we randomly add more screens to the compiled network of interest, it becomes more complete, leading to an increasing overlap fraction from 2% to 5% (Fig. 3a). In contrast, the Normlap scores are less impacted by the incompleteness since the degree inconsistency is already considered, falling within a range of 52–62%, with a slightly decreasing trend as the compiled network becomes more complete. A potential reason for the decreasing Normlap score is that the accumulation of some false positive pairs reduces the network quality as we combine more screens.

**Figure 3.**
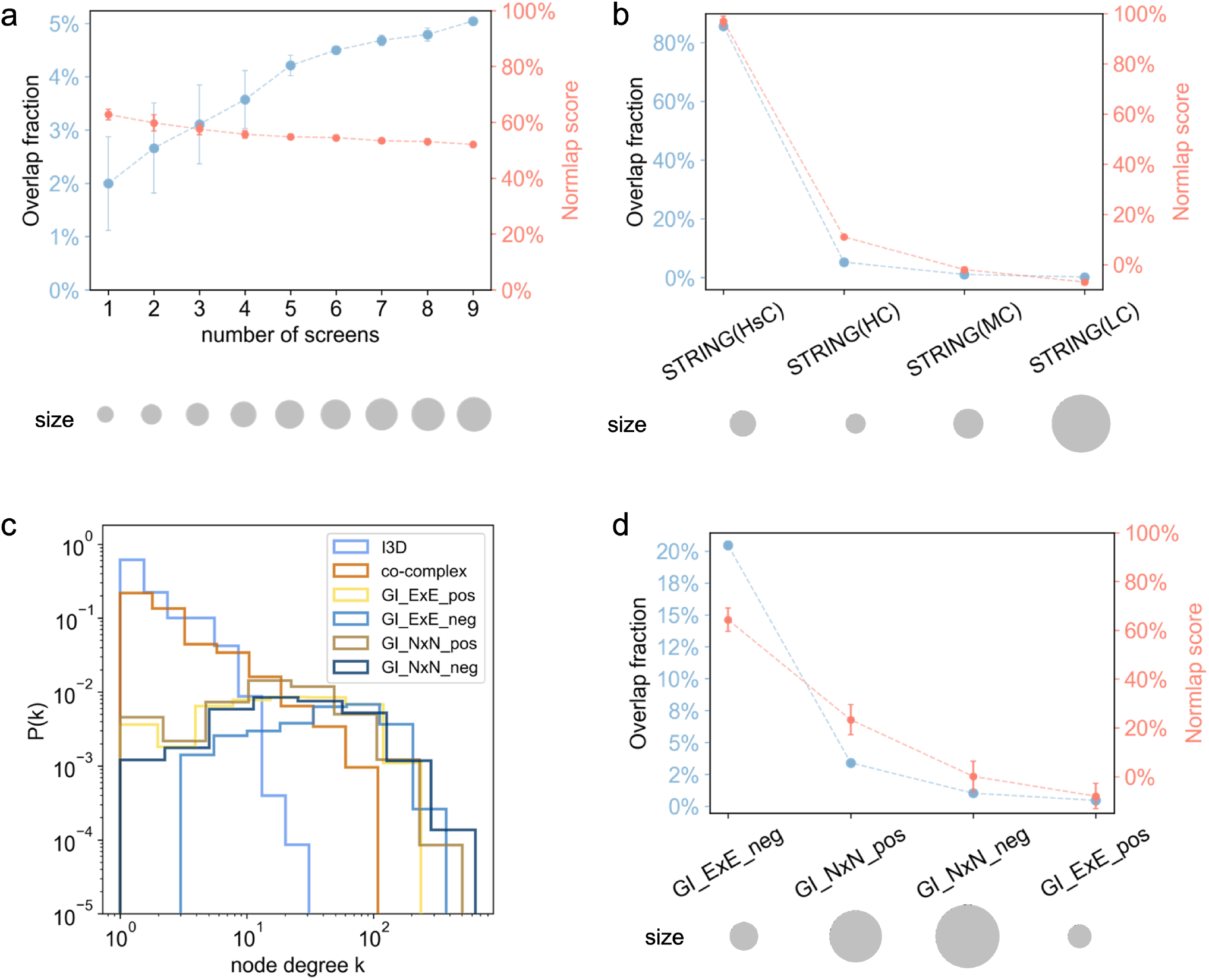
The impact of data incompleteness and quality on the Normlap score. (a) Normlap score and overlap fraction as the number of considered screens increases from the HuRI experiments. The screens are randomly selected. The number of links in each network is indicated by the size of the gray disks. (b) Overlap fraction and Normlap score between I3D and parts of STRING with different confidence score thresholds. (c) Node degree distribution for various yeast networks. (d) Normlap score between the I3D network and GI networks. ExE denotes the interactions between essential genes and NxN denotes interactions between non-essential genes.

Next, we used yeast datasets to further investigate how the Normlap score changes with network quality, by comparing I3D as a gold standard to subsets of the STRING dataset of varying confidence scores. We divided PPIs in STRING into four networks according to the confidence score, forming STRING(HsC), STRING(HC), STRING(MC), and STRING(LC) in decreasing order of network quality (see Methods for more details). As shown in Fig. 3b, both the overlap fraction and the Normlap score decrease as the network quality becomes lower. However, since a lower overlap fraction could either come from decreased network completeness or network quality, no conclusion about network quality can be drawn from the overlap fraction alone. In contrast, the decreasing Normlap scores are capable of indicating the decreasing network quality regardless of the degree inconsistency caused by network incompleteness and other contributing factors.

At a broader scope, Fig. 1c shows the Normlap scores between the gold standard I3D and ten yeast PPI datasets along with the corresponding negative and positive benchmarks, as well as the observed overlaps. All yeast PPI networks scored higher than 89%, except STRING(LC) and RRS. Although the observed overlaps of systematic networks are lower than other networks, the systematic networks are of close quality to other networks, as the observed overlaps are not significantly lower than the positive benchmark except for Tarassov and YeRI.

Comparison of PPI networks to functional networks plays a vital role in understanding the biological rules of a given organism^25^. However, the amplified degree inconsistency (Fig. 3c) between PPI networks and functional networks usually leads to a slight overlap at best, hindering further interpretations. As a starting point, in yeast, there is a unique genome-wide systematic functional network of genetic interactions (GI)^13,14^ proven to be highly successful in mapping the wiring diagram of cellular function. Genetic interactions occur between pairs of genes whose simultaneous mutations enhance (positive interactions) or suppress (negative interactions) the fitness compared to the expectation based on the fitness of single mutants. Previous studies^13,14^ have shown that negative interactions among essential genes (genes critical for an organism’s survival) significantly overlap with PPIs and the co-complex network. This significant overlap is neither observed for non-essential genes nor for positive interactions between essential genes. In addition, positive interactions between non-essential genes overlap with PPIs, albeit to a lesser extent^14^.

Here, as an example in yeast, we have utilized the Normlap score to compare the GI networks^14^ with five PPI networks and the co-complex network^26,27^ (Fig. 3d, S2a-f). Although the overlap fraction of negative interactions between essential genes (GI_ExE_neg) with the other six networks varies from 7% to 32%, the Normlap scores are consistently around 60%. This consistency indicates that a large fraction of the GI_ExE_neg network shares compatible biology to PPI and co-complex networks, in line with previous observations^13,14^. For non-essential genes, the overlap fraction of negative interactions (GI_NxN_neg) ranges from 2% to 7%, while the Normlap scores are consistently around 20% except for Lit-BM-20 (11±1%). The consistent Normlap scores indicate that a lesser fraction of the GI_NxN_neg network shares compatible biology with the PPI and co-complex networks compared to the GI_ExE_neg network. Overall, in the example of yeast and human networks, we showed that the Normlap score is robust against data incompleteness and reflects the agreement of networks reliably.

### Agreement network of networks

The positive benchmark enables us to construct agreement networks even among networks of different sizes and functions. In the agreement network, networks are connected if compatible, i.e. when the hypothesis of having a best-case scenario cannot be discarded at a confidence level of *p* = 0.05. Fig. 4a,c shows the Normlap scores between various networks for yeast and human, followed by the visualization of the agreement networks in Fig. 4b,d.

**Figure 4.**
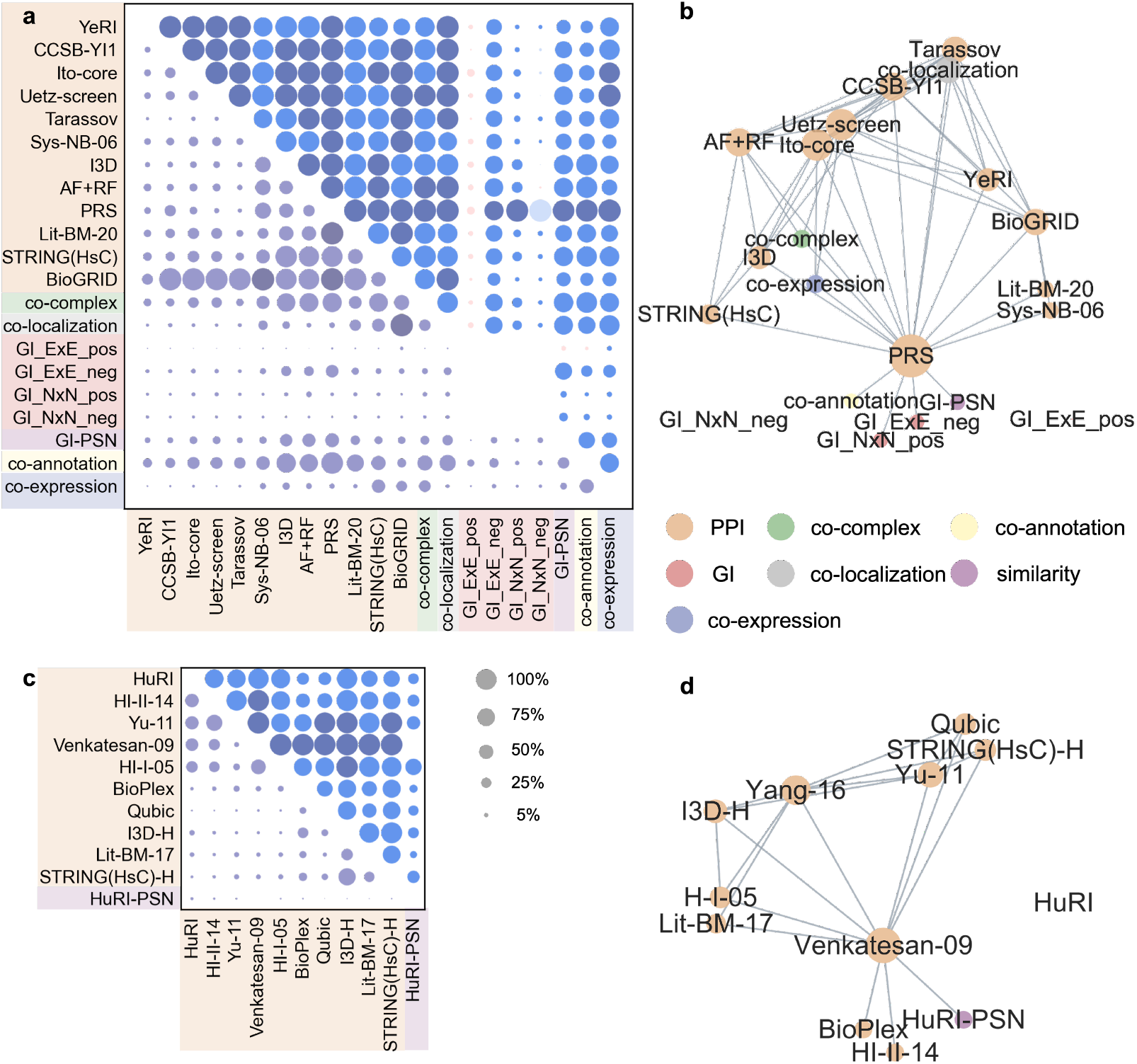
Agreement network of networks. Normlap score (upper triangle) and observed overlap fraction (lower triangle) between (a) yeast and (c) human networks. Negative Normlap scores are colored in red. Pairs with no significant signal are shown in a lighter color. Compatible network pairs (*p* ≥ 0.05) are shown in a darker color. Agreement network of (b) yeast and (d) human networks. Nodes are connected if compatible, see text. The node size is proportional to the node degree.

Note that the Normlap score needs to be interpreted with caution when applied to the case where one network is a subset or almost a subset of the other network. This is because the Normlap score relies on the overlap to infer the quality of the rest of the network. If one network is a subnetwork of the other network, the score will always be 100%, indicating that the subnetwork is consistent with the larger network, which is true, but not informative of the quality of the rest of the links in the larger network. For example, CCSB-YI1, Ito-core, Uetz-screen, Tarassov, Sys-NB-06, PRS and Lit-BM-20 are all subnetworks (or nearly subnetworks) of BioGRID, thus leading to 100% Normlap scores (Fig. 4a). Apart from the special case of BioGRID, all PPI network pairs gained a Normlap score higher than 69%, showing a tendency of consistency.

The difference between binary and non-binary experimental methods can be seen in Fig. 4a. Five binary PPI networks, namely YeRI, CCSB-YI1, Ito-core, Uetz-screen and Tarassov, are compatible with each other. In contrast, these binary PPI networks are not compatible with the non-binary PPI network Sys-NB-06. Since protein pairs in Sys-NB-06 are not necessarily in direct contact, the Sys-NB-06 network has a lower Normlap score (69±4%, *p* < 0.05) with the co-localization network, compared to that between binary networks and the co-localization network (86–100%, *p* ≥0.05). Besides, the PPI networks and the co-complex network are not compatible except for PRS, Uetz-screen and AF+RF networks, which reflects the fact that PPIs do not only happen within complexes but also between and outside of complexes. YeRI has a slightly lower Normlap score (76 ± 4%) compared to other systematic PPI networks (> 89%) when compared to the co-complex network. The lower Normlap score likely originates from the fact that there are more PPIs in YeRI that occur between or outside of complexes, as pointed out in Ref. [6]. Interestingly, we found that structure-based PPI networks I3D and AF+RF have higher Normlap scores (> 84%) than other systematic PPI networks (27–53%) when compared to the co-annotation network. The high Normlap scores indicate that the genes of proteins with compatible binding interfaces are more likely to have functional associations compared to the corresponding gene pairs of other PPIs when the node degrees are taken into account as a confounding factor.

For human networks, all four Y2H binary networks show a high level of consistency with the lowest Normlap score at 73±3% between HuRI and Yu-11. The Normlap score can be much lower when comparing human networks of different experimental methods. For example, the lowest Normlap score of systematic networks is between HuRI and BioPlex (31 ± 1%, *p* < 0.05), indicating there are significant differences between HuRI and BioPlex. The discrepancy may be explained by the fact that Bioplex has significant biases compared to HuRI, which has a more homogeneous coverage^5,17^. Overall, both agreement networks for yeast and human are well connected, highlighting a high level of consistency between networks of the same organism. Indeed, we can connect most of the studied network datasets to others either directly or indirectly, through pairs of compatible networks.

### Threshold and validate experimental assays

Suppose we have a candidate network where the links are ranked by an intrinsic candidate score. The positive benchmark enables us to threshold such scored networks, making them compatible with a gold standard network. Indeed, we can compare the candidate network constituted by the ranked links above a selected threshold to a gold standard network and select the threshold where the filtered network becomes compatible with the gold standard. As an illustration, Fig. 5a,b shows the yeast GI-PSN network ranked by PCC scores and the human Qubic network ranked by enrichment scores, respectively. Precision is calculated as *TP/D*, where *TP* is the overlap and *D* is the total number of links with candidate scores above the threshold. The yeast GI-PSN network is compared to two gold standard networks, I3D and co-complex. We found that the Normlap scores remain consistent for top-ranked links, while the precision varies substantially depending on the selected gold standard networks. The selected GI-PSN network with PCC = 0.50 (corresponding to 962 links) is compatible with the co-complex network, while I3D gives a threshold at 0.55 (corresponding to 458 links). Turning now to the human Qubic network, which is compared to I3D-H and BioPlex as gold standard networks, the selected Qubic network is compatible with I3D-H at *enrichment score* = 11.5 (382 links) and compatible with BioPlex at *enrichment score* = 13.5 (52 links). Therefore, we can systematically threshold candidate networks so that they become compatible with one or multiple gold-standard datasets.

**Figure 5.**
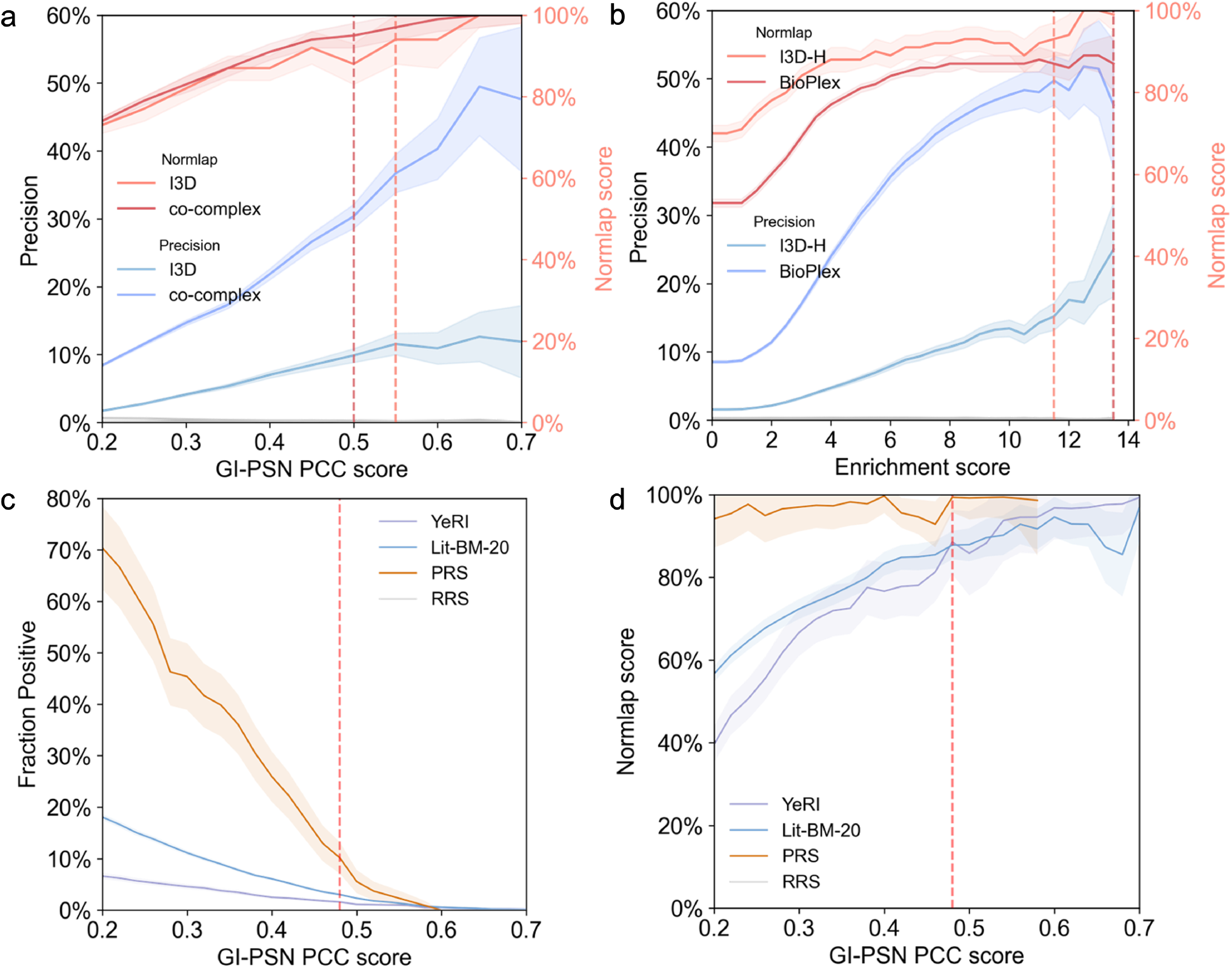
Network evaluation with Normlap. Validation of (a) the yeast GI-PSN with I3D and co-complex networks and (b) the human Qubic with the I3D-H and BioPlex networks. The precision is given by the true positives divided by the total positive predictions. Shades indicate the error bars from shot noise, while gray lines indicate the random reference. (c)-(d) Results of computational validation of YeRI in GI-PSN. Lit-BM-20, PRS and RRS are shown for reference. Red dashed lines indicate the suggested thresholds, see text for details.

In addition to thresholding scored candidate networks, the positive benchmark allows a computational alternative to validate experimental assays. Traditionally, the quality of new experimental datasets is routinely validated by retesting promising links in complementary assays^4–6^. For example, the Y2H v4 screens of YeRI are retested in two complementary binary PPI assays MAPPIT^20^ and GPCA^21^, alongside Lit-BM, PRS and RRS as shown in Extended figure 2b in Ref. [6]. In these examples, the new dataset is considered high-quality if it has a close positive fraction compared to PRS or Lit-BM at the threshold where the recovery rate of RRS was zero. While retesting with MAPPIT and GPCA assays provides a way to validate the experimental datasets, it comes with sizeable costs to retest batches of the experimental data in different labs. Besides, the positive fraction is expected to depend sensitively on the degree inconsistency and whether the tested links are within complexes or outside of complexes (Fig. 2b,c in Ref. [6]), making it difficult to assess the validation results. Furthermore, due to the high cost, there is currently no MAPPIT or GPCA network to systematically assess and address the degree inconsistency.

As a computational alternative, we propose to validate new experimental datasets with an existing scored network. First, the links in the new datasets are ranked by the scores in the existing network. Then, we generate the positive benchmark from the union of the new dataset and pairs above a selected threshold in the scored network. Last, we choose the threshold where the new dataset is compatible with the scored network at the selected threshold. As proof of concept, we validated YeRI with the GI-PSN network (Fig. 5c,d), along with Lit-BM-20, PRS and RRS for reference. Links in the four networks are ranked by the PCC scores in the GI-PSN network, and at PCC = 0.48, YeRI is compatible with the GI-PSN network. Just like in the experimental validation with MAPPIT or GPCA assays, we have comparable normalized positive fractions of the validated network YeRI (89 ± 8%) with Lit-BM-20 (88 ± 3%) and PRS (99 ± 4%) while RRS being 0% at the selected threshold, serving as validation of the YeRI methodology. Note that we could choose a lower threshold as long as the RRS is sufficiently low (0%).

## 3 Discussion

Complex biological networks require multiple complementary approaches to be fully characterized, leading to datasets of low overlap. With the number of assay versions increasing, it becomes more and more crucial to achieve proper quality control of the generated datasets. The quality of datasets from different approaches is often benchmarked by the observed overlap with a gold standard network. However, the observed overlap is not a reliable measure of network quality due to multiple factors contributing to the degree inconsistency. In this study, we proposed a positive statistical benchmark as a generative model for the best-case scenario to assess network quality. The positive benchmark together with the negative benchmark leads to our normalized network overlap score - Normlap.

We demonstrated that the Normlap score is robust to data incompleteness and capable of reflecting network agreement, with the example of human and yeast networks. Furthermore, the positive benchmark and Normlap score enable a standardized comparison of different molecular and functional networks. As a case study, we evaluated the agreement between various yeast and human network datasets. In contrast to the low observed overlap, we observed a widespread consistency among networks of the same organism, as most datasets form a cluster of compatible networks. Although we used I3D and I3D-H as gold standard networks, other networks may serve as gold standards if they are compatible with most existing networks, for example, the human Venkatesan-09 network. Just like there is no single superior assay to map biological networks^3^, there might not be a universal gold standard network either. Yet, our work can be an important step toward creating the next generation of PRS and RRS datasets. Last, we showed a computational alternative to threshold and validate experimental assays using the positive benchmark and Normlap score, instead of experimentally retesting existing networks.

A limitation of the proposed positive benchmark is that it is overestimated for the following reasons. First, we take the union of the two networks as a proxy for the ideal network, while the ideal network can be more complete, leading to a smaller overlap. As a potential solution, the union can be made more complete to gain a better estimate of the true underlying network. Instead of taking the union of two networks, we can add more networks to the union if it is reasonable to assume these networks originate from the same underlying biology. Second, all the links in the union are assumed to be true positives while the union likely includes some noise. The impact of possible noise can be illustrated by considering the extreme case where one network is randomized. Even in this case, the positive benchmark will be higher than the negative benchmark (Fig. S1b), meaning that treating all links in the union as true positives, in this case, leads to an overestimated positive benchmark. More generally, the Normlap score depends nonlinearly on the amount of noise in the datasets. Therefore, additional steps are needed to reliably assess the noise level of the networks, through a process of systematic calibration.

In sum, the proposed generative positive benchmark and the associated Normlap score offer a quantitative assessment of network quality, with broad future applications in any areas of measured networks, such as brain^28^ or social networks^29^. We believe that this is a key advancement compared to existing frameworks that are built upon a comparison to a negative benchmark only. The methodology of generating the positive benchmark can be further extended to more complicated networks such as directed, signed or weighted networks^30^, multilayer networks^31^, as well as dynamical networks^32^. Moreover, the generative model of the positive benchmark allows comparison of other aspects of the two networks, such as distance between networks^33^, with the best-case scenario.

## Methods

### Network construction

Yeast gene names have been mapped to the ORFs (open reading frames) with the SGD database^34^. Genes that could not be mapped to nuclear ORFs have been excluded from our analysis. For consistency among datasets, we have removed the hyphens in the ORFs so that ORF like “YDR374W-A” became “YDR374WA”. Human datasets were mapped via gene or protein identifiers to the Ensembl gene ID space with the hORFeome Database 9.1^35^. Genes that cannot be mapped to Ensemble gene ID are removed from this analysis. All self-interactions in yeast and human datasets are excluded from this analysis. Summaries of the number of nodes and links for all yeast and human datasets used are shown in Table S1 and Table S2.

#### Yeast PPI networks

##### Systematic Y2H PPI networks - YeRI, CCSB-YI1, Ito-core, Uetz-screen

YeRI is the latest systematic map constructed using a novel Y2H assay version after testing ~ 99% of the yeast proteome^6^ three times with assay Y2H v4. CCSB-YI1^4^ is an earlier proteome-scale dataset of Y2H PPIs validated using the two complementary assays MAPPIT^20^ and GPCA^21^. Ito-core^7^ is a subset of PPIs found three times or more in Ito et al.^7^, excluding unreliable pairs of proteins found only once or twice. Uetz-screen^6^ is a subset of PPIs from Uetz et al.^8^ that was obtained from a proteome-scale systematic Y2H screen, excluding a smaller-scale, relatively biased, targeted experiment with a smaller number of well-studied bait proteins.

##### Physically-proximal PPI network - Tarassov

The Tarassov network is a proteome-scale dataset generated using a dihydrofolate reductase protein-fragment complementation assay (DHFR PCA) by Tarassov et al^12^. It contains physically proximal, but not necessarily directly-contacting protein pairs.

##### Systematic AP-MS PPI network - Sys-NB-06

The Sys-NB-06 dataset is obtained from Ref. [6], which contains PPIs from three AP-MS experiments, namely Gavin 2002, Gavin 2006 and Krogan 2006.

##### Inferred PPI network with experimental structures - I3D

The I3D dataset used in this analysis is a subset of Interactome3D^36^ released in May 2020. The subset is restricted to experimental structures, with interactions from homology models excluded. Proteins with compatible binding interfaces are identified as PPIs in I3D.

##### Predicted PPI network from structures - AF+RF

AlphaFold (AF)+RoseTTAFold (RF) is a deep-learning-based predicted PPI network, downloaded from Ref. [6]. The subset used in this analysis is restricted to links with PPI score ≥ 0.9.

##### Literature curated datasets - Lit-BM-20

The Lit-BM-20 dataset is from Ref. [6]. Links with two or more pieces of evidence including at least one binary evidence were selected as Lit-BM-20 links.

##### Positive reference set (PRS) and Random reference set (RRS)

The PRS and RRS dataset is from Lambourne et al.^6^.

##### BioGRID

The BioGRID yeast PPI dataset is constructed from the 4.4.210 release of the BioGRID database^37^. The Organism ID and Experimental System Type are set to be “559292” and “physical”.

##### STRING

The STRING yeast PPI network is constructed from the v11.5 release STRING database^23^. The physical subnetwork is segmented into four networks with different confidence levels, namely highest confidence network STRING(HsC) (quality score ≥ 900), high confidence network STRING(HC) (700 ≤ quality score < 900), medium confidence network STRING(MC) (400 ≤ quality score < 700) and low confidence network STRING(LC) (150 ≤ quality score < 400).

#### Yeast functional networks

##### Yeast genetic interaction (GI) networks

Genetic interaction networks are constructed from the SGA data by Costanzo et al.^14^. The source data is used at the intermediate threshold (*P* < 0.05 and genetic interaction score |*ε*| > 0.08). For interactions with multiple *P*-value and *ε*, the *ε* corresponding to the smallest *P* is used for classification. The same process is applied to both essential (ExE) and non-essential (NxN) gene interaction source datasets to construct the GI_ExE_pos, GI_ExE_neg, GI_NxN_pos and GI_NxN_neg networks.

##### Yeast genetic similarity network

The genetic similarity networks are constructed from the genetic interaction profile similarity matrices by Costanzo et al.^14^. The dataset is filtered with PCC > 0.2 to construct the genetic similarity network (GI-PSN).

##### Yeast co-complex network

The yeast co-complex network was generated from the list of protein complexes from Baryshnikova 2010^26^ and Benschop 2010^27^. Genes in the same complex are connected. Note that 495 genes are classified into multiple complexes, leading to bridges between complexes.

##### Yeast co-localization network

The yeast co-localization network is constructed from the BioGRID^37^ database (4.4.210 release). The Organism ID and Experimental System are set to “559292” and “co-localization” to filter for yeast co-localization links.

##### Yeast co-annotation network

Genes are considered co-annotated if they share annotation from the non-redundant set of specific GO terms^38^.

##### Yeast co-expression network

The co-expression data are from https://coxpresdb.jp^39^. The union dataset Sce-u.v21 was used. Union coexpression is calculated by the average of the logit-transformed MR values, which is the measure of coexpression strength in COXPRESdb, of RNAseq and microarray coexpression; for gene pairs with only RNAseq coexpression, RNAseq coexpression values were converted to union values by linear regression. Only pairs with scores ranked in the top 5000 were used to generate the co-expression network.

#### Human networks

##### Systematic Y2H PPI networks - HuRI, HI-II-14, Yu-11, Venkatesan-09, HI-I-05

HuRI^5^ is a human ‘all-by-all’ reference interactome map of human binary protein interactions from nine Y2H screenings of ~17,500 × 17,500 proteins. HI-II-14^40^ included binary PPIs from the Y2H screen for interactions within a “Space II” matrix of ~ 13,000 × 13,000 ORFs contained in Human ORFeome v5.1. Yu-11^41^ tested ~ 6,000 × 6,000 ORF search space of human ORFs in the ORFeome 3.1 with Y2H screening. Venkatesan-09^15^ contains high-quality Y2H interactions from four Y2H screens that were performed on a set of ~ 1,800 × 1,800 protein pairs that were initially designed to estimate the coverage and size of the human interactome. HI-I-05^42^ is the first map of the human binary interactome obtained by Y2H screening for direct, binary interactions within a “Space I” matrix of ~ 8,000 × 8,000 ORFs contained in Human ORFeome v1.1.

##### AP-MS PPI network - BioPlex

BioPlex^43^ is a proteome-scale, cell-line-specific interaction network. It results from affinity purification of 10,128 human proteins — half the proteome — in 293T cells. The BioPlex v3.0 data was downloaded from https://bioplex.hms.harvard.edu on February 14th, 2022.

##### Inferred PPI network with experimental structures - I3D-H

The I3D dataset used in this analysis is a subset of Interactome3D^36^ released in May 2020. PPIs are restricted to interactions between human proteins.

##### Qubic

Qubic is a proteome-wide human interactome with the quantitative BAC-GFP interactomics (QUBIC) method^44^. Individual interactions are characterized in three quantitative dimensions that address the statistical significance, interaction stoichiometry, and cellular abundances of interactions.

##### Literature curated dataset - Lit-BM-17

The Lit-BM-17 dataset is from Luck et al.^5^. Each PPI with multiple pieces of evidence, with at least one corresponding to a binary method is annotated as Lit-BM-17.

##### STRING(HsC)-H

The STRING human PPI network is constructed from the v11.5 release STRING database^23^. The network is restricted to the physical subnetwork among human proteins, and to the PPIs with the highest confidence score (≥ 900).

##### Human similarity network - HuRI-PSN

Jaccard similarity (number of shared interaction partners divided by the total number of interaction partners) was calculated for every pair of proteins of degree ≥ 2 in HuRI^5^.

### Construction of positive and negative benchmarks

We consider two graphs *G*(*V*_1_, *E*_1_) and *M*(*V*_2_, *E*_2_). For the positive benchmark, *G* and *M* is combined to form *Q*(*V*_1_ ∪ *V*_2_, *E*_1_ ∪ *E*_2_) that includes all nodes and links from *G* and *M*. The graph ensemble {*Gp*} contains subgraphs of *Q* and is constructed by the maximum entropy approach to preserve the expected value of the node degrees in *G*^22^. For example, to construct *Gp*, we fix the mean subgraph node degree at its original value < *k_i_* >*_Gp_*= *k_i_*(*G*). The resulting probability of having a link between nodes *i* and *j* is given by *p_ij_* = 1/(1 + *α_i_α_j_*), with the expected subgraph degree of a node 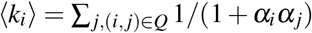. The optimized *α_i_* can be found iteratively, with the update rule 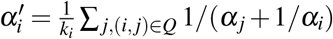. For the negative benchmark, *G* is randomized similarly except that *Q* is replaced by the complete graph of nodes in *G*. We start with the initial condition 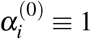 and perform a large number of iterations to update *α_i_*. For the negative benchmark, we stop the iterations when the maximum relative change of *α_i_* is less than 10^−6^. For the positive benchmark, *α_i_* converges slower compared to the negative benchmark due to the high number of constraints. In this case, we stop iterating *α_i_* if the change in the mean overlap of the positive benchmark is less than 1 during the last 1000 iterations. *α_i_* are then used to calculate the connection probability *p_ij_* for all links in *Q*. We can calculate the average and variance of the (positive or negative) benchmark as 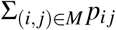 and 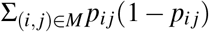, without the need to explicitly generate random samples from the ensemble. This enables an efficient evaluation of statistical significance^22^.

### Visualization of agreement networks

We applied the relative entropy optimization (EntOpt 2.1)^45^ layout to the agreement network in Cytoscape 3.7.1^46^. In the agreement networks, pairs with the *p* ≥ 0.05 are connected, indicating that the observed overlap is not statistically lower than the positive benchmark. For GI networks between essential genes and non-essential genes, both observed overlaps and positive benchmarks are 0 since they do not have common nodes. We set *p* = 0 when *observed overlap* = *positive benchmark* = 0 so that the corresponding edges will not affect the structure of the agreement network.

## Acknowledgements

We thank L. Lambourne for helpful correspondence and R. Xie, H. S. Ansell, and R. Chepuri for useful discussions.

## Supplementary Materials

**Table S1.**
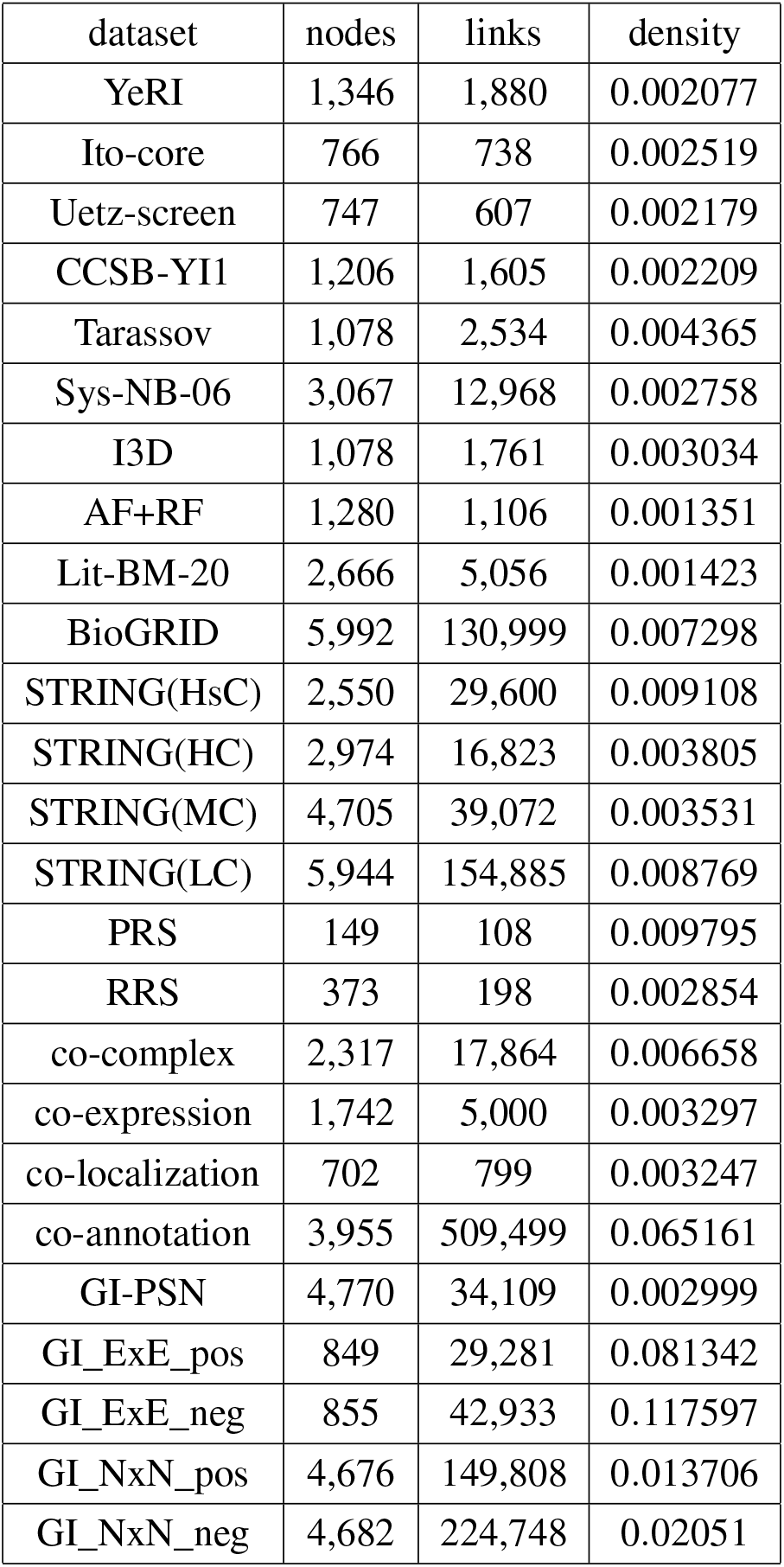
A summary of yeast datasets.

**Table S2.** A summary of human datasets.

| dataset | nodes | links | density |
| --- | --- | --- | --- |
| HuRI | 8,245 | 52,067 | 0.001532 |
| HI-I-05 | 1,500 | 2,561 | 0.002278 |
| Venkatesan-09 | 195 | 187 | 0.009886 |
| HI-II-14 | 4,111 | 13,118 | 0.001553 |
| Yu-11 | 1,133 | 1,127 | 0.001757 |
| I3D-H | 3,155 | 4,354 | 0.000875 |
| BioPlex | 12,408 | 97,078 | 0.001261 |
| Qubic | 4,927 | 23,848 | 0.001965 |
| Lit-BM-17 | 5,956 | 12,758 | 0.000719 |
| STRING(HsC)-H | 6,662 | 39,567 | 0.001783 |
| HuRI-PSN | 4,229 | 33,125 | 0.003705 |

**Figure S1.**
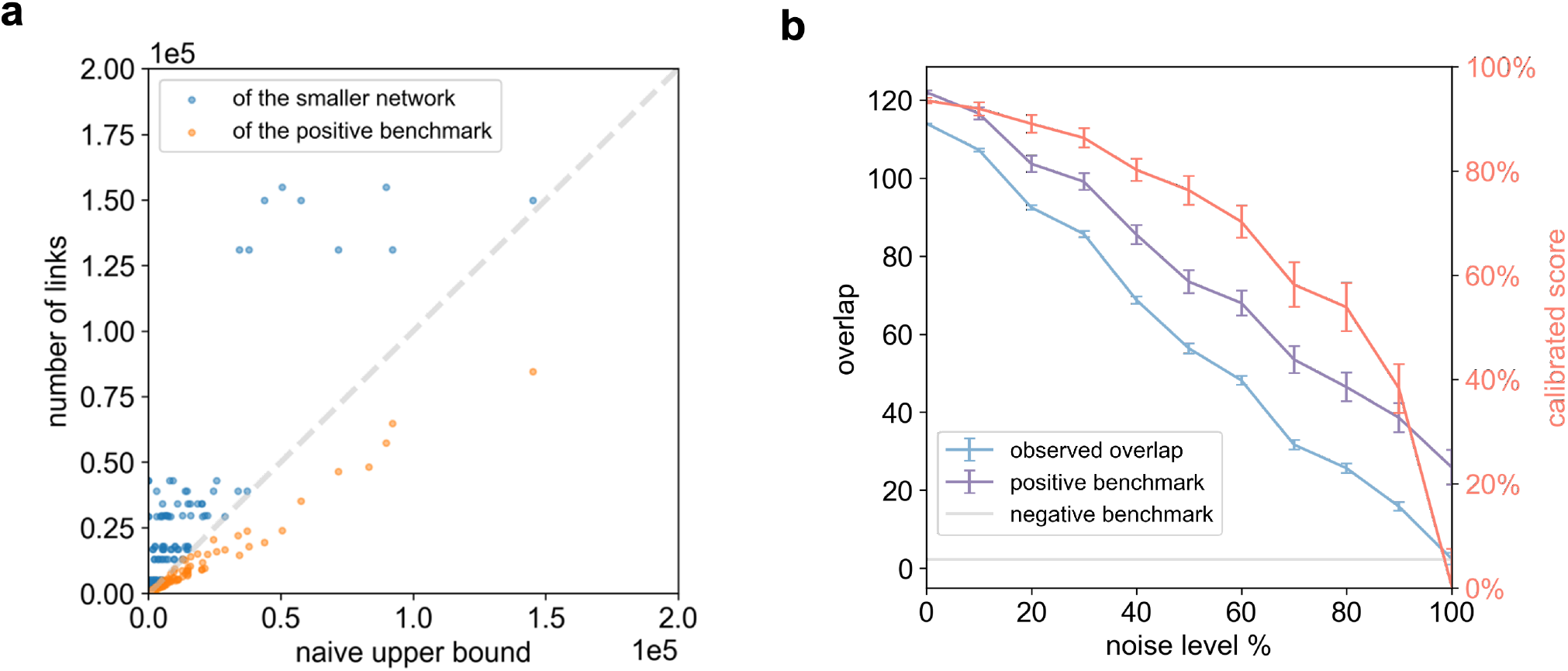
(a) The comparison between the naive upper bound and the number of links of the smaller network and the positive benchmark. (b) Non-linearity of the Normlap score. The CCSB-YI1 is partially randomized according to the noise level while the degree sequence is preserved. The plot showed a comparison with the YeRI network.

**Figure S2.**
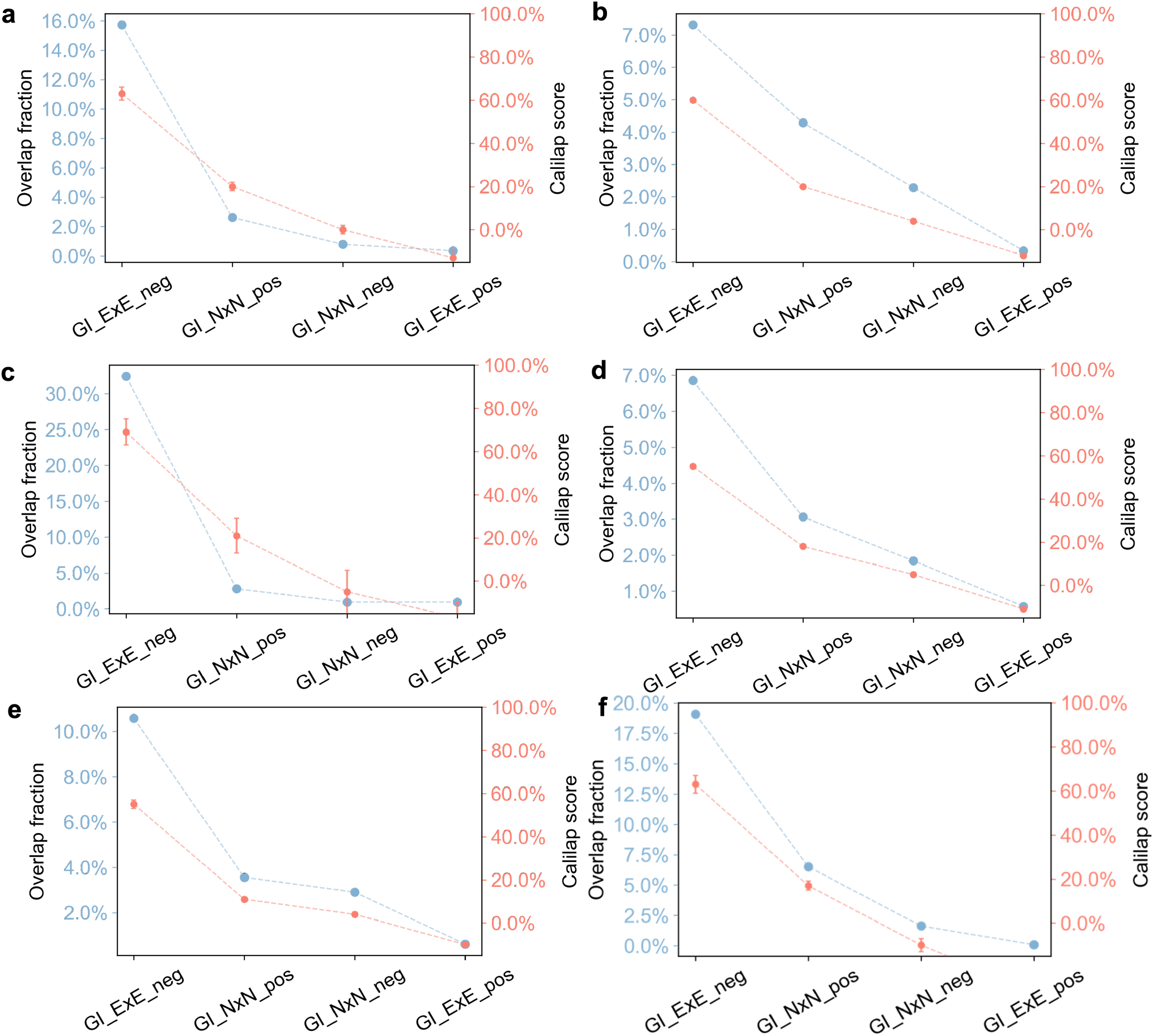
Different aspects of yeast GI network compared with yeast (a) I3D network. (b) co-complex network. (c) PRS network. (d) STRING(HsC) network. (e) Lit-BM-20 network. (f) AF+RF network.

## Notes

### Competing Interest Statement

The authors have declared no competing interest.

